# Loricarioid catfish evolved skin denticles that recapitulate teeth at the structural, developmental, and genetic levels

**DOI:** 10.1101/2021.05.17.444419

**Authors:** Carlos J. Rivera-Rivera, Nadezhda I. Guevara-Delgadillo, Ilham A. Bahechar, Claire A. Shea, Juan I. Montoya-Burgos

## Abstract

The first vertebrate mineralized skeleton was an external bony armor coated with dental structures. The subsequent emergence of a mineralized endoskeleton and of teeth are considered key innovations in the diversification of vertebrates. Although time clouds our understanding of the initial evolution of these mineralized structures, recent re-emergences may shed light on the underlying processes. Loricarioid catfishes are a lineage that, much like the ancestral vertebrates, bear denticle-clad bony armor from head to tail. Loricarioid denticles (LDs) and oral teeth are very similar in superstructure. We show here that other extra-oral dental structures are found as ancestral characters only in lineages that are distantly related to loricarioids such as sharks or coelacanth, indicating that LDs have independently re-emerged in loricarioid catfishes. We investigate whether the similarities between LDs and teeth extend to their developmental and genetic context, and how their development compares to that of other vertebrate integument structures. Our detailed study of the development of LDs, and gene expression analyses through *in situ* hybridization confirm that all 12 genes from the tooth-forming gene regulatory network (oGRN) are expressed in developing LDs in a similar way as they are expressed in developing teeth. We then compare the developmental, structural, and genetic aspects of LD and teeth with that of other integument appendages such as fish scales, shark dermal denticles, feathers and hairs. We find that LDs share all developmental cues with teeth and, to a lesser extent, with the other vertebrate integument structures. Taken together, our results indicate that denticles have re-emerged on the trunk of loricarioid catfishes through the ectopic co-option of the oGRN rather than the resurrection of an ancestral trunk-specific denticle genetic pathway.

## Introduction

Denticulate catfish (suborder Loricarioidei) represent one of a few living lineages of bony fish (Osteichthyes) with dental tissue outside of the oral cavity (Huysseune and Sire, 1997; Rivera-Rivera and Montoya-Burgos, 2017; Schaefer et al., 1989; Schaefer, 1990). These fish are highly specialized to the predator-rich waters of the Neotropics, and the most diverse families (Callichthyidae and Loricariidae) are entirely protected by an armor of bony plates covered with minuscule denticles.

These denticles are often referred to as ‘odontodes’, especially in the morphological and palaeontological literature (i.e. Rivera-Rivera and Montoya-Burgos, 2017; Schaefer and Buitrago-Suárez, 2002; Sire et al., 1998, 1993; Sire and Huysseune, 1996). However, this term could stem confusion in the present work because it is also used in developmental evolutionary biology to refer to a rudimentary developmental structure that is ancestral to all dental and potentially all epithelial structures (Fraser et al., 2010; Ørvig, 1977; Reif, 1982; Stock, 2001). In order to avoid confusion in this work, we will use ‘dental tissue’ to refer to dentin or enamel, ‘tooth’ for mouth skeletal structures bearing dental tissue, ‘denticles’ for skeletal structures bearing dental tissue but located outside the mouth, and ‘placode’ for the developmental (and possibly evolutionary; Fraser et al. 2010) precursor of vertebrate ectodermal appendages, defined as a thickening of the epithelium with a morphogenetic signaling center and communication between the epithelium and the underlying mesenchyme (similar to the interpretation by Pispa and Thesleff (2003).

Throughout vertebrate evolution, denticulate bony armor was not uncommon, and it was most probably the first instance of a mineralized skeleton that emerged in the lineage (Donoghue and Rücklin, 2014; Donoghue and Sansom, 2002). These first denticulate exoskeletons lie at the very heart of the debated question on how and when vertebrates acquired teeth. Historically, the two main hypotheses are: (*i*) the ‘outside-in’ hypothesis, which posits that odontocompetent tissue from the external denticulate body armor folded into the oropharyngeal cavity, leading to the emergence of teeth in jawed vertebrates (Donoghue and Rücklin, 2014; Fraser et al., 2010; Murdock et al., 2013), and (*ii*) the ‘inside-out’ hypothesis, which proposes that teeth first appeared in endodermal tissue within the oropharynx of jawless vertebrates, and then spread onto the external body surface to produce the denticulate bony armor (Fraser et al., 2010; Smith and Coates, 1998; Smith and Hall, 1990). The ‘inside-out’ hypothesis, however, relied strongly on the interpretation that conodonts (ancient organisms with tooth-like structures in their pharynx) are ancestral to all vertebrates. However, a detailed micro-CT study of fossils found that conodonts represent an independent lineage, and their oral tooth-like structures are a convergence with the teeth of jawed vertebrates (Murdock et al., 2013). This study, thus, weakened the ‘inside-out’ hypothesis as a way to explain the emergence of teeth in vertebrates. However, an alternative exists: in their review, Fraser et al. (2010) suggested that teeth are built thanks to an odontogenic gene regulatory network (oGRN), and that dental structures will emerge whenever and wherever this battery of genes (*bmp2, bmp4, ctnnb2, dlx2, eda, edar, fgf3, fgf10, notch2, pitx2, runx2, shh*) is expressed in the presence of odontocompetent neural crest cells (NCCs).

Currently, there are but a few living vertebrate groups that have dental tissue outside their oral cavity (Sire et al., 2009): sharks, skates and rays (Chondrichthyes (Gillis et al., 2017)), coelacanths (Srcopterygii: Actinistia (Smith, 1979)), *Polypterus* (Osteichthyes: Cladistia (Ichiro et al., 2013)), gars (Osteichthyes: Lepisosteiformes (Ichiro et al., 2013)) and denticulate catfish (Osteichthyes: Siluriformes (Rivera-Rivera and Montoya-Burgos, 2017; Schaefer and Buitrago-Suárez, 2002; Sire, 2001)). Body denticles show phylogenetic continuity along the evolution of early jawed vertebrates (Fig. 1), a pattern that remains true when extinct lineages are considered (Donoghue and Rücklin, 2014). However, while extant cartilaginous fish (Chondricthyes) still bear denticles on their trunk, the continuity is broken along the evolution of the two main lineages of bony fish (Osteichthyes). In lobed-finned fish (Sarcopterygii), the phylogenetic continuity of denticles is interrupted in the lineage leading to lungfish (Dipnoi) plus tetrapods (Amphibia plus Amniota). In ray-finned fish (Actinopterygii), body denticles have been lost in two instances: first in chondrosteans (sturgeons and allies – a highly derived group whose members have a naked skin with rows of bony plates), and then at the origin of teleosteans, a group comprising ~35,000 fish species (Fricke et al., 2020), the vast majority of which have elasmoid scales as body cover (Lemopoulos and Montoya-Burgos, 2021), without dental tissue nor denticles (Fig. 1). Within teleosteans, the ostariophysan clade Otophysi (comprising Siluriformes, Cypriniformes, Gymnotiformes, and Characiformes, Nakatani et al., 2011), also contains mostly species with elasmoid scales, with the notable exception of the order Siluriformes (catfishes). The majority of the more than 4,000 catfish species (Fricke et al., 2020) have a naked skin. It is at the origin of the catfish group Loricarioidei (~120 million years ago) that denticles re-emerged in vertebrate evolution (Rivera-Rivera and Montoya-Burgos, 2017). The denticles on the trunk of loricarioids form in close association with underlying bones such as bony fin rays and, when present, dermal bony plates (Rivera-Rivera and Montoya-Burgos, 2017).

**Figure 1.**
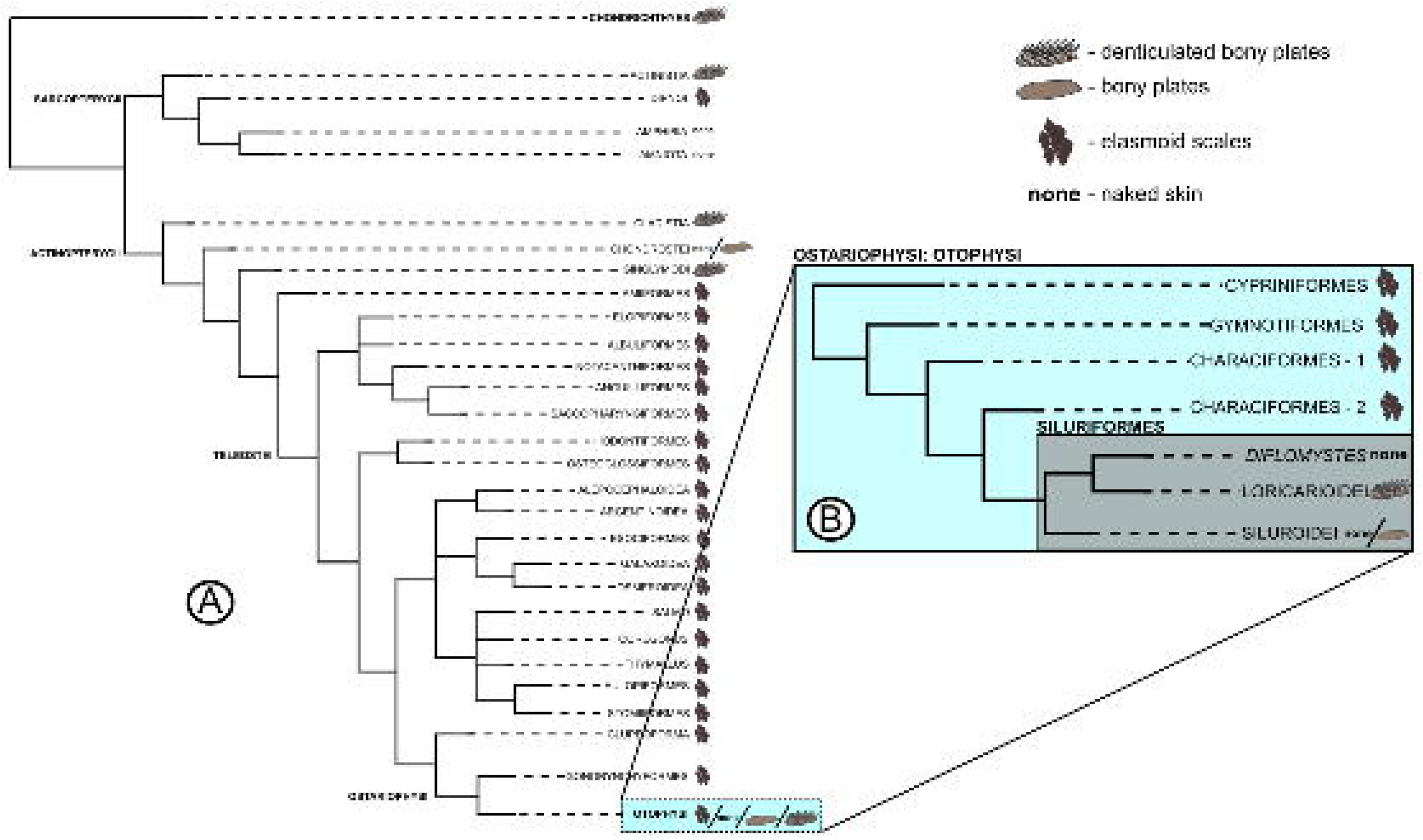
Phylogenetic context of dermal bony elements in Osteichthyes. A phylogeny of Osteichthyes, with Chondrichthyes as an outgroup, with the presence of dermal bony elements shown. In A,, a general overview of the main osteichthyan groups, showing the prevalence of the elasmoid scale, especially within Teleostei. In B., an inset expanding Otophysi, a clade within the superorder Ostariophysi to which the catfish order Siluriformes belongs. In B. also, a breakdown of how the main groups of catfish are related to each other. The phylogenetic data from Osteichthyes was obtained from Diogo (2007), the position of Chondrichthyes relative to Osteichthyes from Donoghue and Rücklin (2014), the branching within Otophysi from Nakatani et al. (2011), and the branching within Siluriformes from Rivera-Rivera and Montoya-Burgos (2017). Characters were obtained for Chondrichtyes from Donoghue and Rücklin (2014), for Dipnoi from Sire et al. (2009), for gars (Ginglymodi) and *Polypterus* (Cladistia) from Ichiro et al. (2013), and for catifsh from Rivera-Rivera and Montoya-Burgos (2017).

Due to the rarity of extra-oral dental tissue in vertebrate species, loricarioids offer a singular opportunity to expand our knowledge of the underlying genetic regulators of integument organs and the morphogenesis of the denticulate armor that was once widely present in early vertebrates. Here, we assess two hypotheses that could explain how LDs were first formed in the loricarioid ancestor. The first is the co-option hypothesis, in which the oGRN deployed to build teeth was co-opted on the body of the loricarioid ancestor through its ectopic activation via changes in the upstream regulatory steps (effectively making LDs ‘teeth outside the mouth’). The second is the resurrection hypothesis, in which an ancestral genetic pathway controlling the formation of dermal dental structures on the body would have been reactivated in the loricarioid ancestor. To assess these hypotheses, we first examined the structural similarities between LDs, teeth shark denticles, and other alternative intergument appendages both as adult structures, and throughout their ontogeny. Second, through gene expression analyses, we determined the similarities between the GRN underlying the development of LD, the oGRN identified for teeth in other vertebrate lineages (Fraser et al., 2009), and putative GRNs activated in the development of the alternative vertebrate integument structures. We used the bristlenose catfish *Ancistrus triradiatus* (Siluriformes: Loricariidae) as a model loricarioid species for the *in situ* hybridization experiments to examine gene expression.

## Results

### Structural similarities between LDs and teeth

We began to address our hypotheses by studying in detail the structure of LDs. A previous study of ours found that adult LDs are histologically almost identical to mature teeth and shark dermal denticles (Rivera-Rivera and Montoya-Burgos, 2017), and a deeper study of the extensive comparative literature on vertebrate dental structures confirms this (see Fig. 2, and the following associated references: Cooper et al., 2017; Martin et al., 2016; Rivera-Rivera and Montoya-Burgos, 2017; Sire, 1990; Sire et al., 2009; Sire and Meunier, 1993). For the present study, we also did histological analyses of developing LD buds, and compared them with the development of teeth, shark denticles, and elasmoid scales (see Fig. 3, and Fig. 4 and associated references: Aman et al., 2018; Cooper et al., 2017; Sire and Akimenko, 2004; Smith and Hall, 1993; Togo et al., 2016). The developmental process of LDs resembles very much that of a tooth and a shark dermal denticle, but does not resemble that of the teleost elasmoid scale (Fig. 4). In particular, while the elasmoid scale forms within the mesenchyme, LDs, as teeth and shark dermal denticles, form in the ectoderm-mesenchyme barrier (see, for example, the cryosections for *notch2* in Fig. 3). This is consistent with the fact that dental structures are ectomesenchymal structures that need an underlying odontocompetent mesenchyme to produce the odontoblasts that deposit dentin, and ectodermal, columnar ameloblasts that deposit enamel on top of that layer (Koussoulakou et al., 2009; Reif, 1982; Sire et al., 2009). In addition, dental structures and elasmoid scales are built of different materials: enamel and dentin for teeth, LDs, and shark denticles vs. elasmodine for scales (Sire, 2001; Sire et al., 2009).

**Figure 2.**
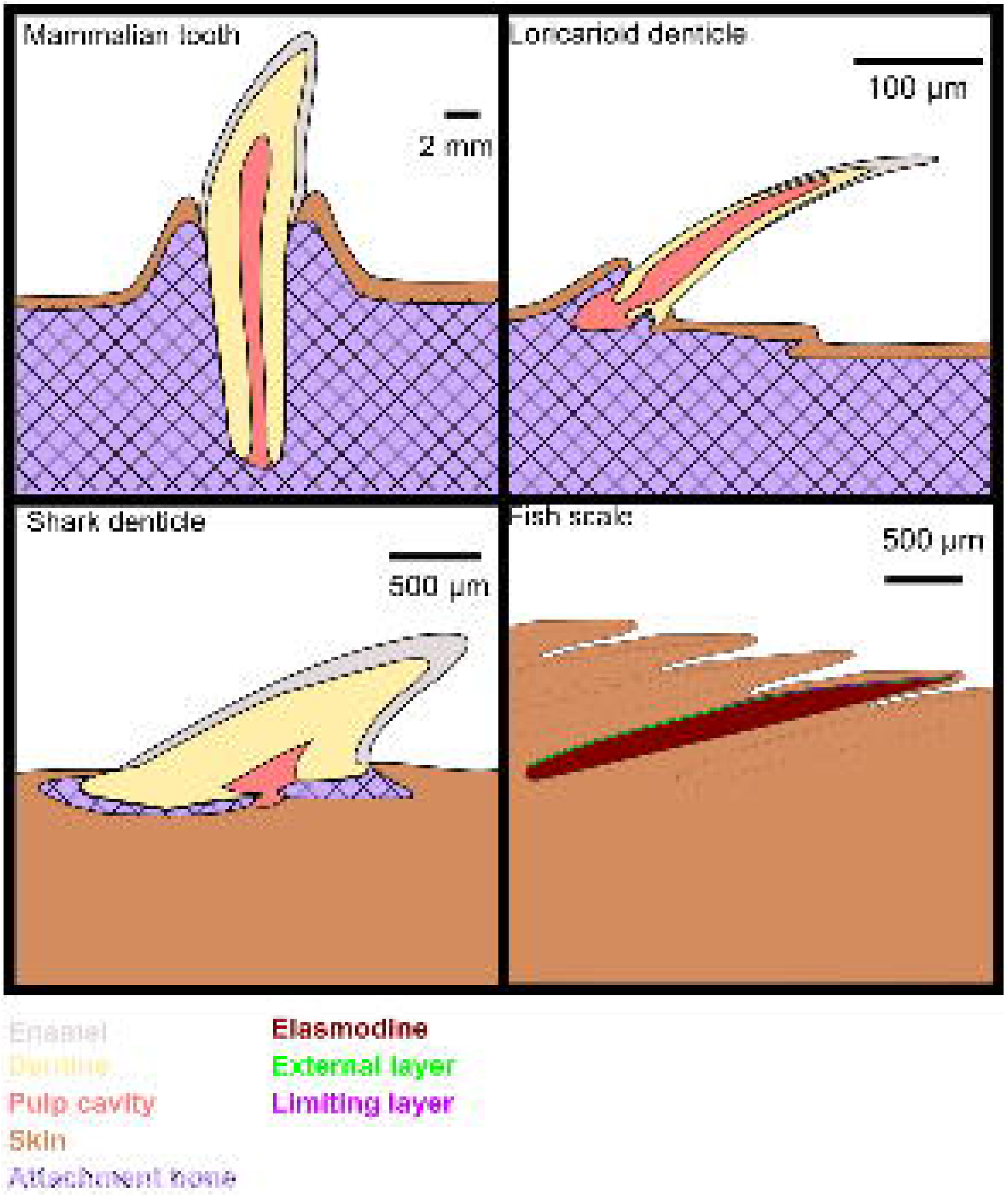
Diagrams of cross-sections of three fully-developed dermal bony structures found in Osteichthyes. Diagrams of cross-sections of a mammalian tooth, a loricarioid denticle, shark dermal denticle, and an elasmoid scale. Histological data come from the following sources: Mammalian tooth, Hall (2015); loricarioid denticle, Rivera-Rivera and Montoya-Burgos (2017) and Sire and Meunier (1993); shark dermal denticle, Sire et al. (2009), and Martin et al. (2016); fish (elasmoid) scale: Sire et al. (2009) and Sire (1990). Note the similarities in tissue organization among teeth, loricarioid denticles and shark denticles, despite the difference in size.

**Figure 3.**
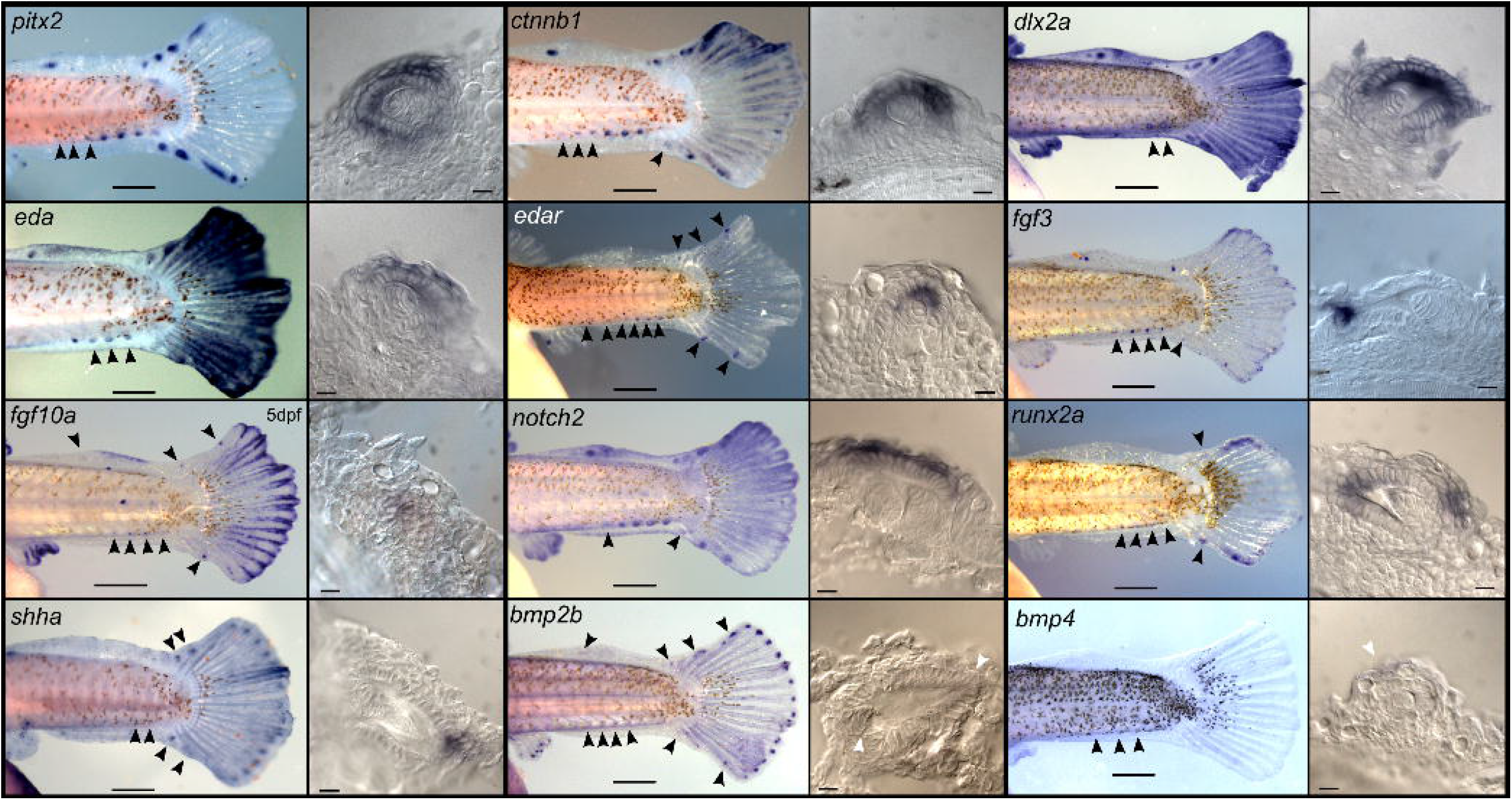
The oGRN deployed in tooth formation is also expressed during loricarioid denticle formation. Gene expression of all 12 genes from the oGRN in caudal denticle buds of *Ancistrus triradiatus* embryos, displaying a punctate pattern associated to denticle buds. Whole-mount ISH are accompanied by histological cryosections showing denticle buds with expression of the gene studied. All embryos, with the exception of the one presented for *fgf10a* are of 6 dpf; *fgf10a* expression is for a 5 dpf embryo. Black arrowheads mark places in which a signal is visible when the image is amplified. Scale bars in the whole-mount images is 500 μm, and 10 μm in the histological cryosections.

**Figure 4.**
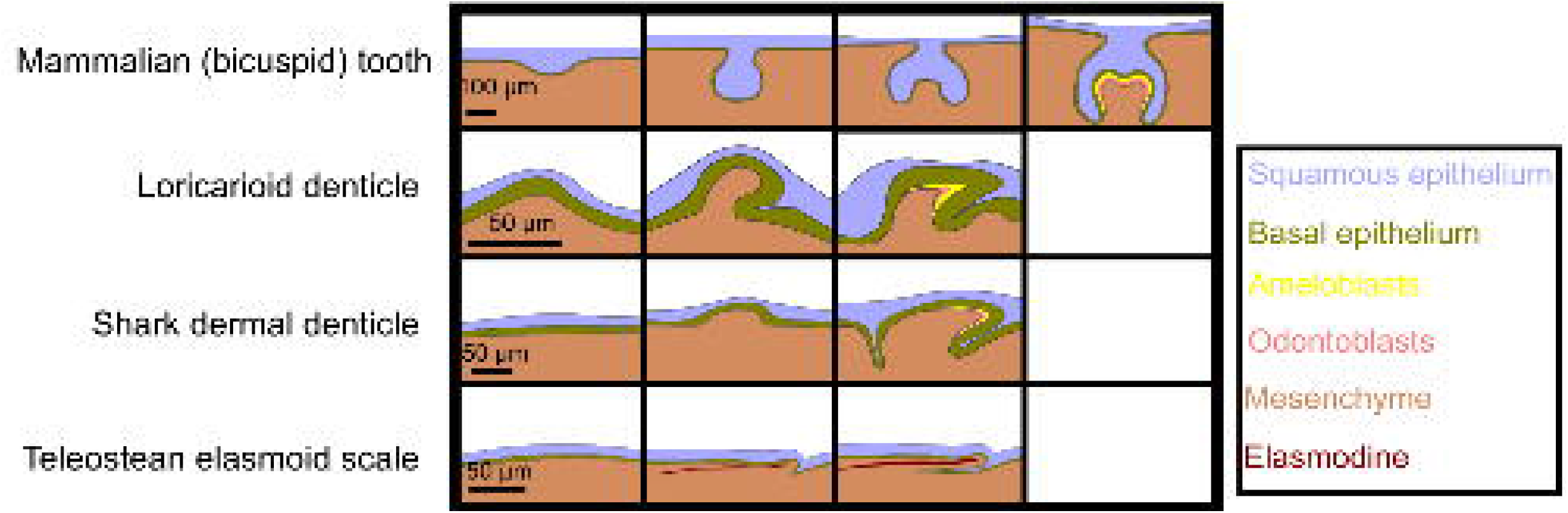
Diagrams of cross-sections of developing dermal bony structures, Details of a developing tooth, a loricarioid denticle, a shark dermal denticle, and an elasmoid scale, with the different ectodermal layers marked. Note how dental structures form at the ectomesenhymal boundary, whereas elasmoid scales form within the mesenchyme. The histological data was collected from the following sources; tooth development: Thesleff (2014), Smith and Hall (1993), Togo et al. (2016); loricarioid denticle: this study; shark dermal denticle: Cooper et al. (2017), and elasmoid scale: Sire and Akimenko (2004), Aman et al. (2018), Sire (1990).

### Determining whether the oGRN is expressed in developing LDs

We then sought to determine whether the genes comprised in the oGRN were expressed in early LD buds in our model species *A. triradiatus*. Being from the Loricariidae family, *A. triradiatus* has one of the most complete denticulate armors in the catfish order, with denticulated bony plates covering its entire body, including the head, the whole trunk (except the abdominal area), and all fin rays. We explored the expression of the oGRN with *in situ* hybridization (ISH) using probes for each of the 12 genes included in this GRN (*bmp2b, bmp4, ctnnb2, dlx2a, eda, edar, fgf3, fgf10a, notch2, pitx2, runx2, shha*) (Fraser et al., 2009). Using alizarin red stains of embryos, we identified that the first LDs emerge on every fin ray, including the adipose fin spine, and on a ventrolateral series, starting at the caudal peduncle (Fig. 5). Thus, we sought to determine whether the expression of oGRN genes co-localized with the places where the first denticles would emerge, on embryos from 2-6 days post-fertilization (dpf). In addition, we did histological cuts of the whole-mount embryos to confirm that the signal observed was associated to a denticle bud, and not any other placodal structure. All 12 genes of the oGRN were expressed both in the mouth and on areas of the trunk where LDs eventually emerge. However, we saw gene expression pattern variation among the different genes studied: the most obvious signal was a punctate expression pattern corresponding to the locations where the first trunk denticles emerge (Fig. 6, Fig. 5): a series along the ventrolateral area of the posterior caudal peduncle, on the first ray of the dorsal, pectoral and adipose fins, and on the dorsal and ventral spines of the caudal fin. This punctate pattern was already visible at 3 dpf for *pitx2, ctnnb1, eda* and *runx2a*; at 4 dpf for *edar, fgf3, notch2, shha* and *bmp2b*; and at 5 dpf for the remaining genes, *dlx2a, fgf10a*, and *bmp4* (Fig Fig. 3 and Supplementary Fig. 1). In the later stages (5-6 dpf), all genes showed this punctate expression pattern (Fig. 3). Some genes were expressed in a smear-like pattern during the earlier stages of development which then turned into a more circumscribed signal on or around putative early denticles (*pitx2, ctnnb1, dlx2a, eda, fgf3* and *notch2*; Supplementary Fig. 1). For the remaining genes, the punctate pattern was the first signal to appear. In addition, we confirmed that each of the oGRN genes was also expressed in developing teeth in *A. triradiatus*, a necessary positive control that ensured that these genes were also being deployed during tooth development, as is the case in many other vertebrates (Supplementary Fig. 2). Histological cryosections on the trunk clearly show that the signal of all 12 genes is associated to denticle buds (Fig. 3). We also noted that the determination of future denticles was not pre-patterned on the skin, as it happens with elasmoid scales (Sire and Akimenko, 2004), but rather seemed to progress as a periodic pattern in which the determination of the first denticle placode begins a wave of successive denticle placode formation that ends up spanning the whole body, similar to was has been observed in the dental lamina of chiclids during the formation of their dentition (Fraser et al., 2008).

**Figure 5.**
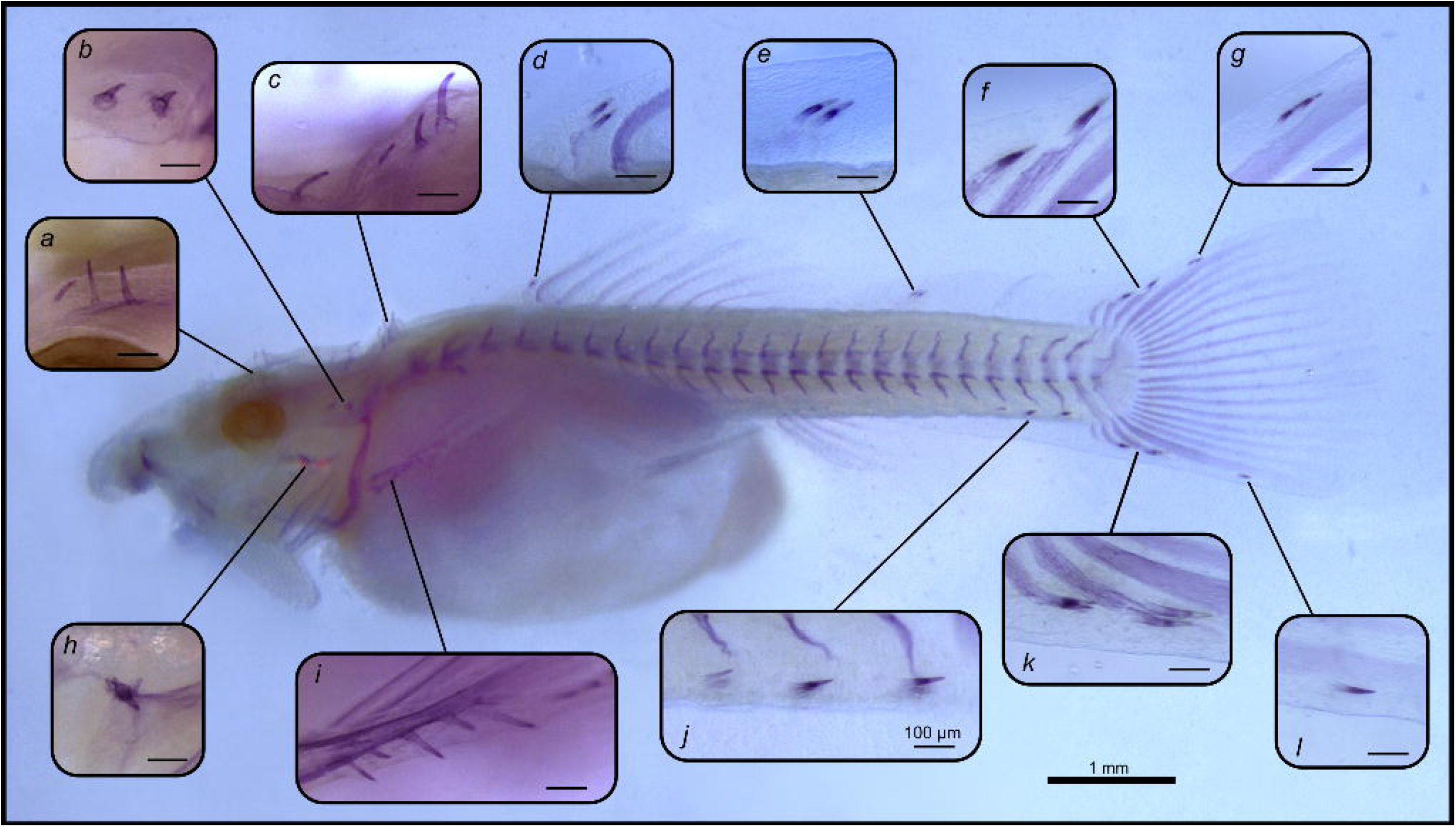
Early denticle mineralization in a 8 dpf embryo of *Ancistrus triradiatus*. Mineralized denticles in an alizarin red-stained 8-dpf *Ancistrus*. (A) Frontal bone denticles, (B) autopteroticum bone denticles, (C) parieto-supraoccipital bone denticles, (D) dorsal fin spine denticles, (E) adipose spine denticles, (F,G) dorsal caudal fin spine denticles, (H) opercular bone denticles, (I) pectoral fin spine denticles, (J) ventrolateral denticle series, (K,L) ventral caudal fin spine denticles. The scale bars in all inset images are of 100 μm, and on the full-body image is 1 mm.

**Figure 6.**
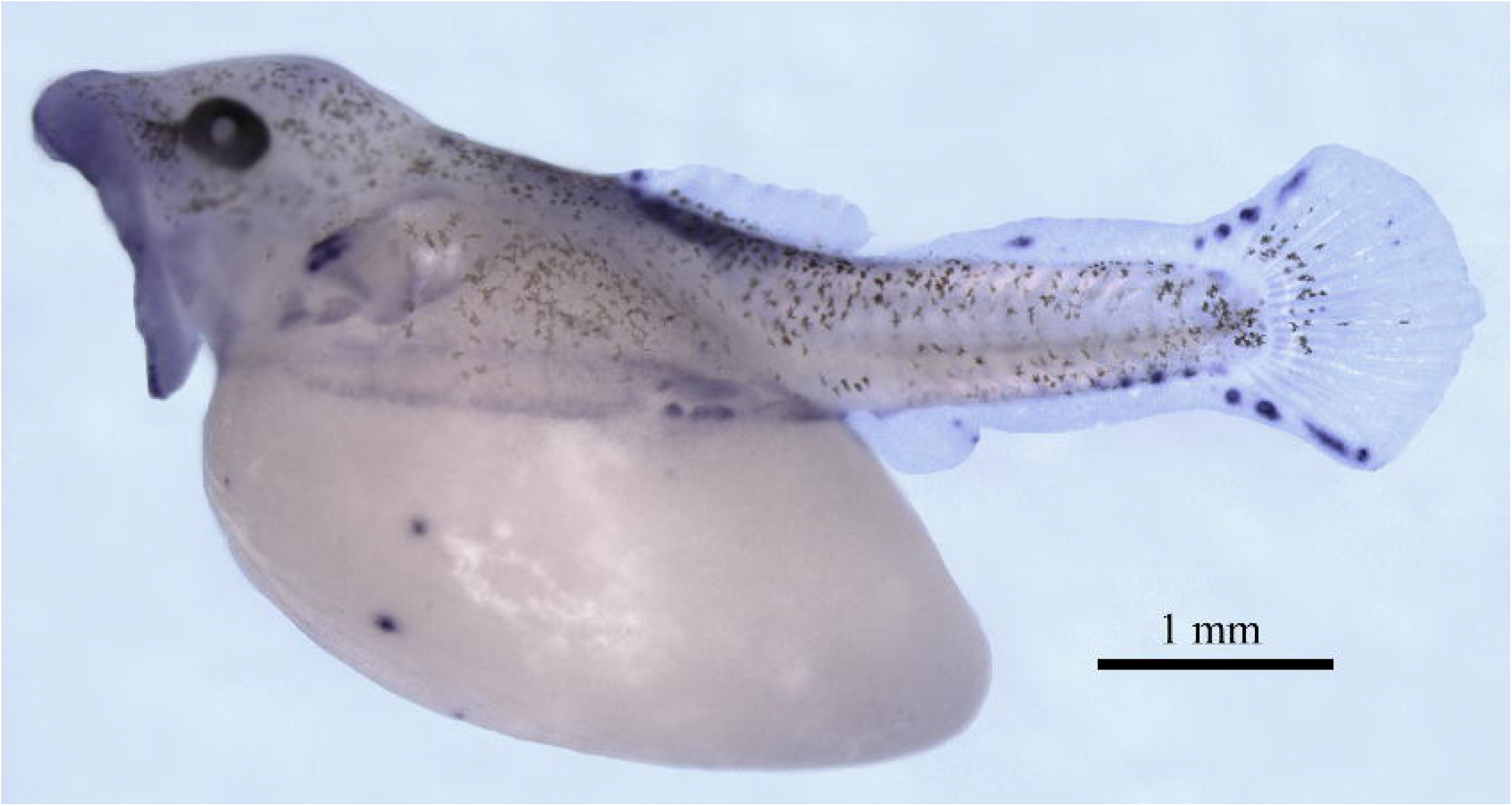
Whole-mount, *in situ* hybridization with a probe for the *pitx2* gene, in a 6 dpf embryo of the armored catfish *Ancistrus triraiatus*. 6 days-old embryo of *A. triradiatus* showing where the tooth-specific gene *pitx2* is expressed on the trunk, prior to denticle formation. The signal is strong in the areas where the first denticles emerge: each fin ray, and a ventrolateral series (corresponding to denticles in Fig. 5 d, e, f, g, i, j, k, l)

### The oGRN genes in LDs, teeth, and other vertebrate integument structures

Based on our results and gene expression data from the literature, we made a presence/absence matrix of all 12 oGRN genes across 17 different integument vertebrate structures, including teeth from several osteichthyan lineages, other epithelial placodal structures and trunk bony structures (Fig. 7). This analysis showed that all 12 genes of the oGRN are found in association to LDs as well as tooth buds within osteichthyan representatives (cichlids, zebrafish, loricarioid catfish, mouse, crocodilians with the exception of the gene *dlx2* which is not found), and in sharks (except for *dlx2*). Interestingly, *pitx2* appears to be a marker of dental placodes, as it is not expressed during the development of all other ectodermal placodal-structures in which its expression has been studied as well as from the two integument bony structures compared (Fig. 7).

**Figure 7.**
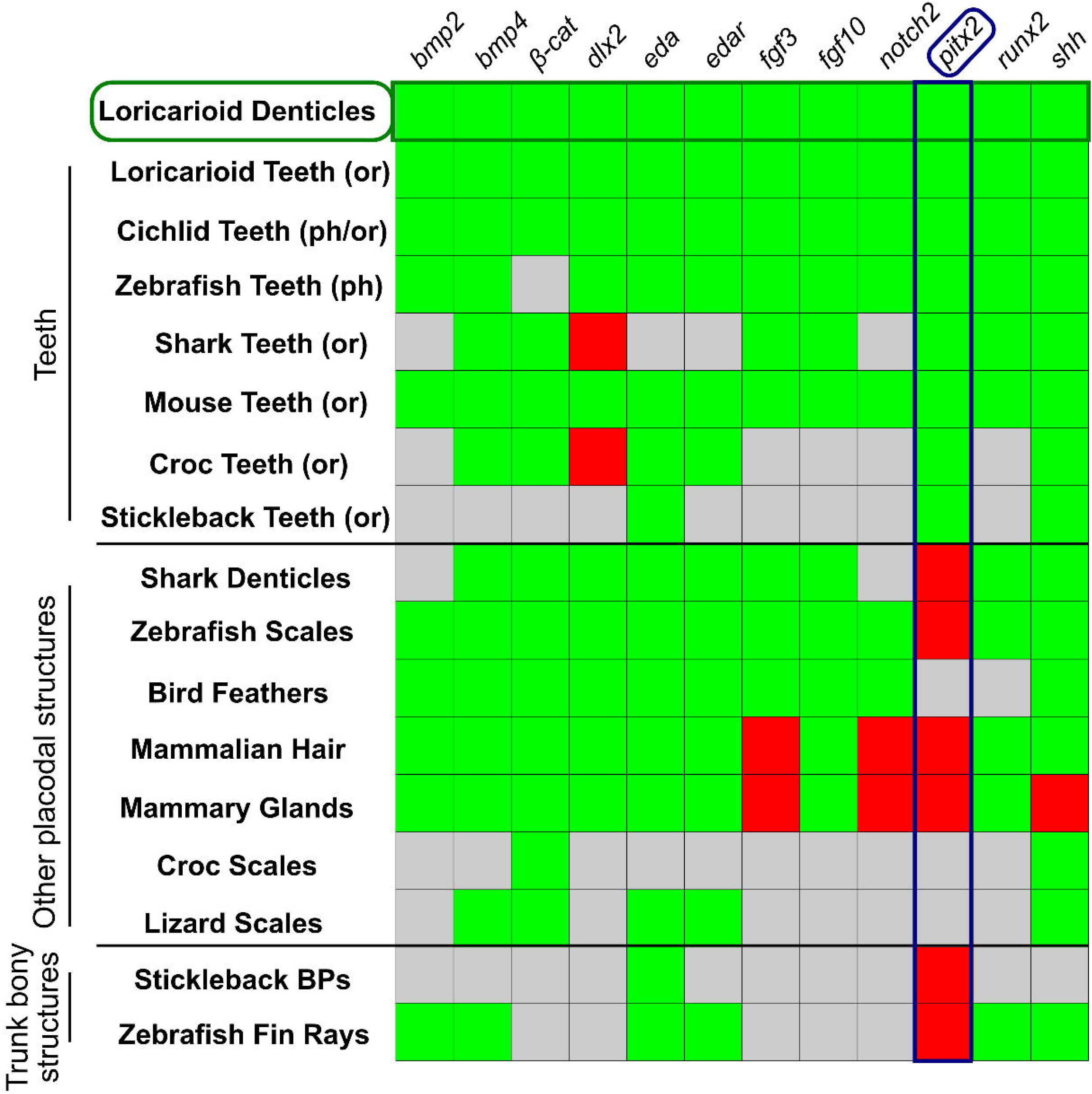
Expression of the 12 genes in the tooth-forming gene regulatory network (oGRN) in 17 different vertebrate bony and placodal structures, Heat map showing expression (green), lack of expression (red), or expression unknown (grey) of each of the 12 genes from the oGRN. The data about loricarioid denticles and loricarioid teeth are from this study, whereas the data for the rest of the table stems from other studies. All 49 references consulted are in the table with the raw data.

## Discussion

The emergence of mineralized dental tissue is a landmark in the evolution of vertebrates (Kawasaki et al., 2004). These hard tissues provided protection as denticles on the trunk, and as teeth in the oral cavity they increased the ability of biting and processing food. Despite the importance of these structures in the evolution of vertebrates, we know remarkably little about the genetic and developmental processes behind their evolution. For example, dental tissue is observed in different places on the body of vertebrates, and was probably gained and lost along the evolution of multiple lineages. It seems possible then, to re-gain a dental structure that was lost ancestrally Yet, are the same GRNs behind their formation, or do they get activated in a different way?

This is, to our knowledge, the first study to test for the expression of the complete battery of oGRN genes in extra-oral dental structures. We find that LDs are very similar to other integument structures containing dental tissue within vertebrates in their superstructure (Fig. 2), development (Fig. 4), and gene expression (Fig. 3). We found *pitx2* as a marker for tooth ectodermal placodes, as its expression is not detected in non-tooth ectodermal placodes, not even in elasmobranch dermal denticles, despite these being structurally very similar to teeth (Fig. 2, Fig. 4, Fig. 7). Based on its early expression in avian teeth of a chick mutant, *pitx2* was already presumed to be an early marker of dental placodes (Harris et al., 2006). Interestingly, we find that the tooth specific *pitx2* gene is expressed in denticles buds during LD formation (Fig. 6, Fig. 3).

The two alternative hypotheses to explain the emergence of dental structures on the body of the Loricarioidei we have put forward are: (1) the co-option hypothesis, which posits that the tooth genetic program was re-activated some place else than in the mouth, also posited by Sire (2001); and (2) the resurrection hypothesis, which suggests that LDs are the result of a re-activated ancestral tooth placodal GRN.

Our results give evidence to support the co-option hypothesis. First, we show that the 12 genes that regulate the development of teeth in *A. triradiatus* are also expressed during the formation of LDs in this same species. Second, the expression of *pitx2*, a marker of tooth placodes, during early LD development is striking genetic evidence supporting the hypothesis that LDs are ‘ectopic teeth’. Moreover, the fact that all members of the full oGRN are found in early LDs indicate that these structures are the result of a recent re-activation of the oGRN, as none of the main regulatory genes are missing. This is in contrast to what is seen in shark denticles, which are presumed to be the direct descendants of the ancient denticulate armor of vertebrates, and lack the expression of *pitx2*. Third, LDs are always found in very close association to some underlying bony structure, such as a bony plate, or highly ossified fin spines (Rivera-Rivera and Montoya-Burgos, 2017) recapitulating the structural association between teeth and jaw bones. Also, both teeth and LDs are linked to their underlying bone through a hinged socket and attachment ligaments (Fig. 2, Rivera-Rivera and Montoya-Burgos, 2017; Sire and Meunier, 1993). These structural commonalities between teeth and LDs are not found in shark denticles or in other integument placodal appendages. In addition (and in contrast to elasmoid scales), LDs do not seem to be pre-patterned, but rather grow in what seems like a periodic pattern, a process already described for developing teeth (Cai et al., 2007; Fraser et al., 2008), feathers (Ho et al., 2019), and hair follicles (Sick et al., 2006) that results from interactions between morphogenetic signals that resemble reaction-diffusion (Turing, 1952) patterns in a specific regulatory background. We mention this because the formation of LDs seems to emanate from ‘seed’ initial denticles that appear, and subsequent denticles begin to form away from the initial one, at a more or less even space.

One question that remains outstanding if LDs are ‘ectopic teeth’ is: what kind of mesenchymal cell line is contributing to LD formation? It is generally assumed that only the cranial neural crest (CNC), which migrates within the head mesenchyme, is capable of differentiating into odontoblasts and thus, produce dentin (Baker, 2008; Donoghue et al., 2008; Hall, 2015). Yet LDs, which contain dentin, are not restricted to the head, and form in association with exposed bones on the trunk. The contribution of trunk neural crest in the formation of denticles is realistic: a recent study of dermal denticle development in a chondrichthyan, the little skate, *Leucoraja erinacea*, found that their dermal denticles were formed with the contribution of the trunk neural crest, not the CNC (Gillis et al., 2017).

Another observation that would lend support to a hypothesis that proposes that LDs are the result of oral developmental signals occurring on the body is that most probably all catfish (Siluriformes) have taste buds covering their body (Dana Ono, 1980). Both teeth and taste buds are oral placode-derived ectodermal organs, and the formation of taste buds on the bodies of catfish may result from an ancestral expansion of normally oral developmental signals into the trunk. This might reflect a preliminary potentialisation step found in all catfish, which enabled the subsequent ectopic activation of the oGRN on the trunk of the loricarioid lineage.

Our results show for the first time that the oGRN is deployed for extra-oral and trunk denticle formation in the same manner as for teeth, and we sought to contextualize this information with what is known on the genetic regulation of other related structures.

## Acknowledgments

We thank Joost Woltering, Isabel Guerreiro, and Leonardo Beccari for help optimizing the whole-mount ISH protocols. We also thank Chloé Hot for her assistance in the production of the transcriptome. This research was funded by the Swiss National Science Foundation (grants #31003A_141233 and #310030_185327 to J.I.M.B), and the Institute for Genetics and Genomics in Geneva (iGE3).

## Competing interests

The authors declare that they do not have any financial or non-financial competing interests.

## Materials and Methods

### Captivity and breeding of fish

Sub-adult and adult specimens of *A. triradiatus* were obtained from the aquarium trade. After a two-week quarantine period, they were moved into growth or breeding tanks. The fish were fed four times a week with cucumber, zucchini, potato, and a protein-enriched vegetable mix made in-house.

Breeding tanks were kept at 26° C under a 12-hour light/dark cycle, with a gravel or sand bottom with a piece of wood and bricks which served as potential places for egg laying to take place visibly. Once one or several males had taken residence in available places, we changed large volumes of aquarium water (~20%) every other day, refilling with chilled deionized water. This simulates the flow of water that occurs during the tropical wet season and promotes reproduction. Overnight, one or two females would lay eggs in a male’s hiding spot, and the eggs were subsequently fertilized and looked after by this male for approximately the next ten days.

### Sample fixation

All embryos were collected directly from the aquarium in which they were laid, at the desired age (2-6 dpf). Unhatched embryos (younger than 5 dpf) were dechorionated by hand, with forceps. All embryos were euthanized in a tank containing a lethal concentration (400g/ml) of tricaine methanesulfonate (MS-222), then fixed in 4% paraformaldehyde (PFA) pH 7.42 for at least 24 h at 4° C. Then, they were dehydrated in a series of methanol solutions and stored in 100% methanol at −20° C.

Juveniles were freshly collected upon death due to natural causes. Throughout the first weeks of growth, there are several ‘checkpoints’ at which large-scale deaths are observed, especially during the transition from yolk-sac feeding to eating (7-9 dpf, which also coincides with the time that the first denticles begin to mineralize), granting the opportunity to collect older specimens without having to euthanize them. Once they were collected, they were fixed in 4% PFA pH 7.42 for 24-48h and 4° C, depending on the animal’s size, and then progressively dehydrated to 80% ethanol in distilled water for storage at −20° C.

### *Alizarin red staining and clearing of* A. triradiatus *specimens*

Fixed *A. triradiatus* specimens were first placed in ethanol 100% overnight, and then rehydrated in distilled water through a decreasing series of ethanol concentrations. In the last three steps of rehydration, KOH was also added in increasing concentrations (0.02%, 0.05%, 0.1%). Pigmentation was then removed using a solution of 3% H_2_O_2_ / 0.1% KOH in distilled water, and then the embryos were transferred into the alizarin red stain solution for 10 min. To produce this solution, 10 μl of stock alizarin red solution (1 g Alizarin Red S from Acros Organics, 400460100 in 100 ml 0.5% KOH) were added per 1 ml of 1% KOH. Then, a final clearing step was done with 1% KOH, and for storage and image acquisition, they were placed in 100% glycerol anhydrous overnight, or until the specimen sank.

### *Sequencing and assembly of the embryonic transcriptome of* A. triradiatus

In order to design probes for ISH we needed the sequences of the expressed transcripts that we wanted to target, so we sequenced and assembled the embryonic transcriptome of *A. triradiatus*. Two embryos of 3, 4, and 5 dpf were collected, pooled and their total RNA was extracted using the Trizol reagent. The mRNA was then purified using the Oligotex mRNA Mini Kit (Qiagen 70022). Library preparation, quality control and Illumina NGS sequencing was performed by the Genomics Platform of the University of Geneva (http://www.ige3.unige.ch/genomics-platform.php). Sequencing was performed on a HiSeq 2500 (Illumina) which generated 60 million single reads of 200bp. Illumina adapters and low quality bases were trimmed from reads using Cutadapt v1.9.1 (Martin and Wang, 2011). Read quality was checked using FastQC v0.11.5 (http://www.bioinformatics.babraham.ac.uk/projects/fastqc/). *De novo* assembly was performed using Velvet v1.2.01 (Zerbino and Birney, 2008), with multiple k-mer lengths (k-mers: 35, 45, 55, 65, 75), following the protocol suggested by Surget-Groba and Montoya-Burgos (2010). Multiple k-mers assemblies were then merged using the merge module implemented in Oases with a k-mer length of 27. The raw sequencing data will be made available on NCBI’s Sequence Read Archive (SRA).

### ISH RNA probe design and synthesis

A transcript for each gene was located in our transcriptome by using BLAST+ v2.2.28 (Camacho et al., 2009) to find the corresponding gene sequence of zebrafish (*Danio rerio*) as obtained from the ENSEMBL database. After identifying each gene transcript, a 400-700 bp long section was selected as the probe sequence. When possible, approximately half of the probe’s length lay in either the 5’ or the 3’ untranslated regions (UTR), in order to increase signal specificity. Once the probe sequence was selected, we designed primers (Supplementary Table 1) to amplify that section from cDNA using a standard polymerase chain reaction (PCR) and identified the successful amplifications through agarose gel electrophoresis.

Well-amplified fragments were cloned into a pGEM-T vector using the pGEM-T Easy Vector System (Promega A1360), following the manufacturer’s instructions and bacteria were cultured on LB-agar plates supplemented with ampicillin (150 ug/ml), X-gal (100 ul/plate) and IPTG (10 ul/plate). After an overnight incubation at 37° C, the white colonies were collected; half of each colony was transferred into a new culture plate, and the other half was lysed in 50 μl of NP40 0.1% for PCR. Successful bacterial transformation was confirmed by PCR with Dir2/Rev2 standard primers and orientation of the DNA insert was determined by PCR with Dir2 and our designed forward primers. The Dir2/Rev2 PCR products were used for Sanger sequencing using the standard primer pair M13F/M13R (Fasteris, Geneva, Switzerland). For probe synthesis, we selected plasmids whose DNA insert had no mutations from the DNA used for transformation (if there were none, we chose the sequence with the highest fidelity), and was oriented such that SP6 could be used to create an anti-sense probe (in our experience, SP6 has been more effective than T7 in generating our RNA probes).

The probe template was amplified with a standard PCR reaction using the insert’s forward primer and Rev2 or Dir2 as the reverse primer (depending on insert orientation; the promoter to be used must be included in the amplified fragment). In most cases, several PCR reactions had to be combined and purified in a smaller volume in order to reach the minimum fragment concentration of 1 μg/13 μl needed for probe synthesis.

The probes were synthesized using the DIG RNA Labeling Kit (Roche 11175025910) following the manufacturer’s instructions. The RNA probe was then purified with the RNAEasy Kit (Quiagen, 74104), following the “RNA cleanup protocol”, and eluting in 35μl of RNAse-free water. To visualize synthesized probe quality, 1μl of the probe was run on a fresh 1.7% agarose gel. Probe concentration was determined using a Nanodrop ND-1000 and stored at −80° C.

### *Optimized whole-mount ISH protocol for* A. triradiatus *embryos*

Our ISH protocol is a modification of a previously published protocol for ISH of non-model vertebrates (Woltering et al., 2009), and has aspects from a second protocol optimized for fish (Thisse and Thisse, 2008). Unless otherwise stated, all of the following steps were done on a shaking platform at room temperature.

Embryos collected and fixed as described above were first bleached in a solution of 1.5% H_2_O_2_ in methanol for at least 30 min or more, depending on the amount of pigmentation. We also performed this step for early embryos with little or no pigmentation, as we found that it improved the penetration of the probe. The embryos were then rinsed in 100% methanol, and rehydrated for 10 min in 50% methanol in millipore-filtered water, 10 min in TBS-T (15 mM NaCl, 2.5 mM KCl, 25 mM Tris base, 0.1% Tween-20, pH 7.4), and twice for 5 min in TBS-T.

The embryos were permeabilized in a fresh solution of Proteinase K (Roche 02115887001) diluted to 10 μg/ml in TBS-T. The time of Proteinase K treatment varied with embryo size, and we found that *A. triradiatus* embryos of 3 and 4 dpf needed 8 min, 5 dpf ones 10 min, and 6 dpf ones 10 min. Next, the embryos were placed in 0.2 M triethanolamine (Sigma-Aldrich T58300) pH 7-8, to which 25 μl of acetic anhydride per ml of triethanolamine was added. The embryos were kept in this solution for 20 minutes or more until the acetic anhydride drops had dissolved completely. After three 5 min washes in TBS-T, the embryos were re-fixed in 4% paraformaldehyde in PBS (137 mM NaCl, 2.68 mM KCl, 101 mM Na_2_HPO_4_, 1.76 mM KH_2_PO_4_, pH 7.4) for 20 min, and then washed three times for 5 min in TBS-T.

All hybridization steps took place at 68° C, and all solutions were preheated before applying them to the embryos. The embryos were prehybridized in Hybmix A (250 μl/ml of 20x SSC made with 3 M NaCl, and 341 mM C_6_H_5_Na_3_O_7_ at pH 7.5, 50% deionized formamide, 50 μg/ml heparin, 0.3 mg/ml torula RNA, 200 μl/ml of a 4% stock solution of blocking reagent (Roche 11096176001) which was dissolved and autoclaved in MNT solution (150 mM maleic acid pH 7.5, 300 mM NaCl, 0.1% Tween-20), 10 mM EDTA, and 2% Tween-20) for 4 h, and then incubated overnight in Hybmix A with the probe at a concentration of 100 ng/ml. The following day, the embryos were washed twice for 1 h in Hybmix B (5x SSC pH 7.5, 50% deionized formamide, and 2% Tween-20), and four times for 30 min in 2x SSC with 0.2% Tween-20.

At room temperature, the embryos were washed two times for 5 min in TBS-T, and then placed in Blocking buffer (120 μl/ml of the 4% stock solution of blocking reagent dissolved in MNT, and 10% heat inactivated goat serum) for 1 h. Embryos were incubated in anti-DIG-AP (Roche 11093274910) antibody (1:4000) in Blocking Buffer for 5 h. Following antibody incubation, the embryos were washed in MNT by first doing a quick wash, then a 5 min wash, a 10 min wash, two 20 min washes, and finally by leaving the embryo overnight in MNT. The next day, hourly washes in MNT were done, and the embryos were again left overnight in MNT. Extending this step is possible if a large amount of nonspecific antibody binding is suspected.

When staining was ready to begin, the embryos were first washed 3 times for 10 min in TBS-T, then 2 times for 10 min in AP-Buffer (0.1 M Tris base pH 9.5, 0.1 M NaCl, 1 mM MgCl2, 0.2% Tween-20). Finally, the embryos were incubated in pure BM Purple (Roche 11442074001) in the dark, from 1 h to one or two days, depending on the speed of color development. The embryo coloration was monitored closely in this step. Once the desired coloration was achieved, the embryos were washed three times for 10 min in AP-Buffer, six times for 10 min in TBS-T, and left overnight in TBS-T. The next day, the embryos were washed twice for 5 min in TBS-T, then re-fixed in 4% PFA in PBS (pH 7.4) for 2 h. After three more washes in TBS-T, the embryos were photographed in TBS-T with a Leica M205C or a Leica MZ16FA with a fluorescence module.

For some embryos, DAPI staining was done in order to have a better image of the body surface of the specimen. This was done by immersing the embryos in 0.1 mM DAPI (Sigma-Aldrich D9564) in TBS-T for 1 h, shielded from light, and then washing them in TBS-T twice for 15 min.

All whole-mount ISH experiments were repeated three times to ensure that the observed signal was replicable.

### Cryosectioning and imaging of whole-mount ISH

After whole-mount ISH, embryos for which more detailed information on the localization of gene expression was needed were prepared for cryosectioning by soaking them overnight in a solution of 30% sucrose in PBS. Cutting blocks were then prepared on dry ice by pouring optimal cutting temperature compound (OCT, TissueTek) into moulds made from aluminum foil and placing the samples within them. Once completed, the cutting blocks were left overnight at −80° C. Cryosectioning was done on a Leica CM1850, and 8 μm sections were placed onto SuperFrost/Plus slides (Assistent), and immediately covered in a solution of polyvinyl alcohol (Mowiol, Sigma-Aldrich) and a glass cover slip, and left overnight to dry. Imaging was done using Nomarski interference contrast with a Zeiss Axioplan 2 microscope fitted with a Leica DFC300 FX camera.

## Notes

### Competing Interest Statement

The authors have declared no competing interest.

