## Supplementary figures and images for "Loricarioid catfish evolved skin denticles that recapitulate teeth at the structural, developmental, and genetic levels"

### Supplementary Figure 1

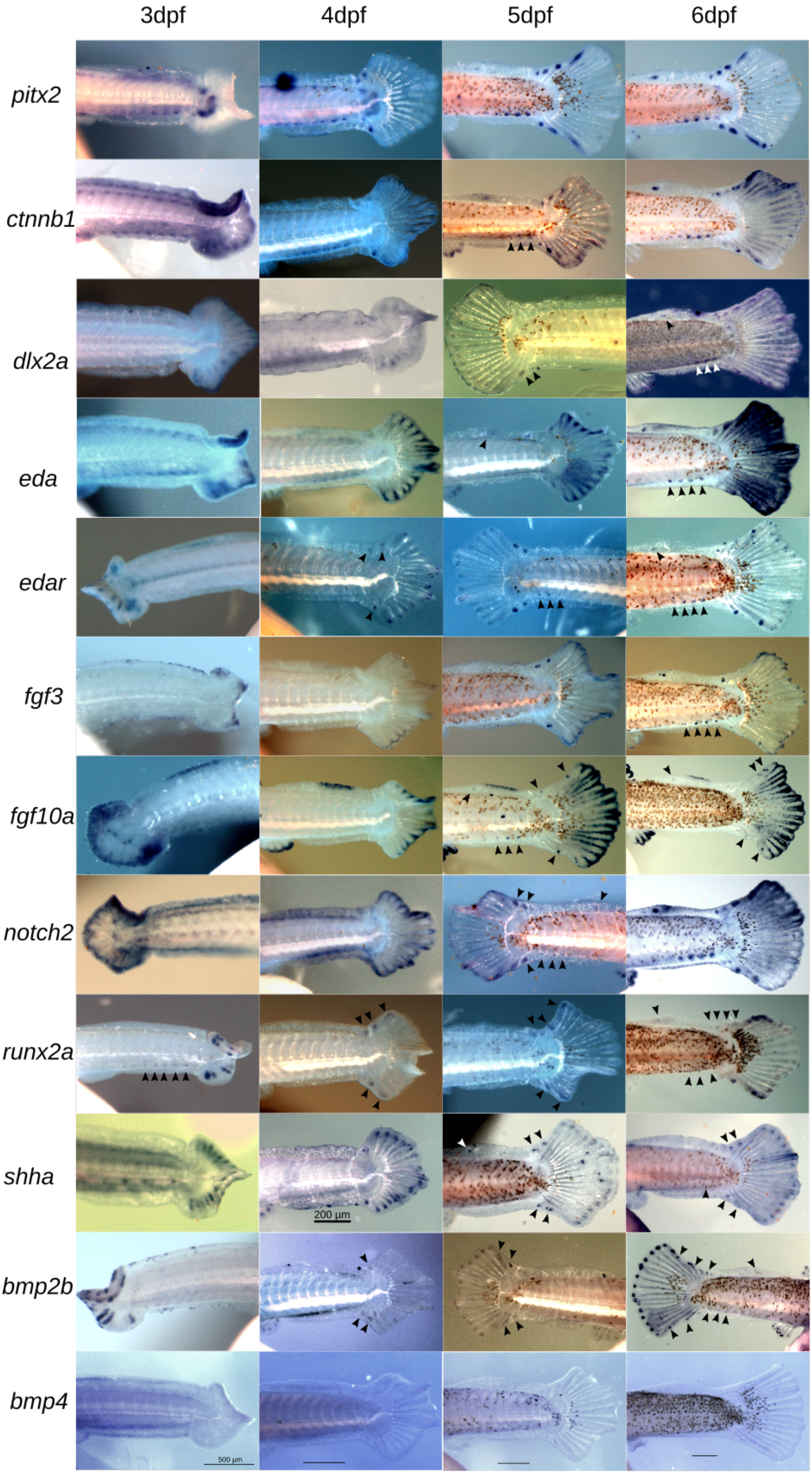

### Supplementary Figure 2

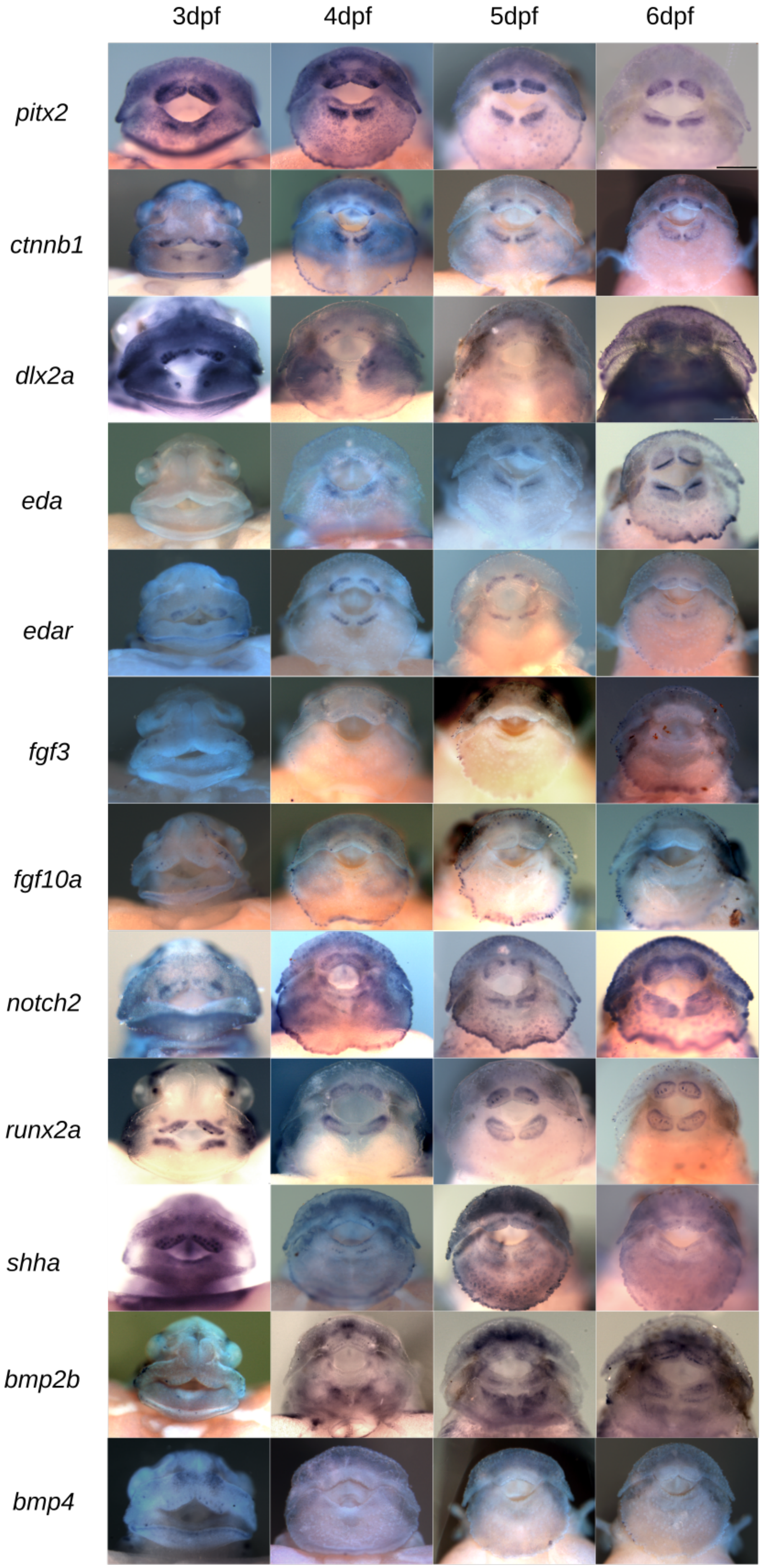
